# Trait-based approach to bacterial growth efficiency

**DOI:** 10.1101/427161

**Authors:** Mario E. Muscarella, Xia Meng Howey, Jay T. Lennon

## Abstract

Bacterial growth efficiency (BGE) is the proportion of assimilated carbon that is converted into biomass and reflects the balance between growth and energetic demands. Often measured as an aggregate property of the community, BGE is highly variable within and across ecosystems. To understand this variation, we first identified how species identity and resource type affect BGE using 20 bacterial isolates belonging to the phylum Proteobacteria that were enriched from north temperate lakes. Using a trait-based approach that incorporated genomic and phenotypic information, we characterized the metabolism of each isolate and tested for predicted trade-offs between growth rate and efficiency. A substantial amount of variation in BGE could be explained at both broad (i.e., order, 20 %) and fine (i.e., strain, 58 %) taxonomic levels. While resource type was a relatively weak predictor across species, it explained > 60 % of the variation in BGE within a given species. Furthermore, a metabolic trade-off (between maximum growth rate and efficiency) and genomic features revealed that BGE is a predictable metabolic feature. Our study suggests that genomic and phylogenetic information may help predict aggregate microbial community functions like BGE and the fate of carbon in ecosystems.

**Originality and Significance:** Bacterial growth efficiency (BGE) is an important yet notoriously variable measure of metabolism that has proven difficult to predict. To better understand how assimilated carbon is allocated, we explored growth efficiency across a collection of bacteria strains using a trait-based approach. Specifically, we measured respiration and biomass formation rates for populations grown in minimal media containing one of three carbon resources. In addition, we collected a suite of physiological traits to describe each strain, and we sequenced the genome of each organism. Our results suggest that species identity and resource type may contribute to growth efficiency when measured as an aggregate property of a natural community. In addition, we identified genomic pathways that are associated with elevated BGE. The findings have implications for integrating microbial metabolism from the cellular to ecosystem scale.

## Introduction

In most ecosystems, heterotrophic bacteria play a pivotal role in determining whether organic carbon is respired as carbon dioxide (CO_2_) or converted into biomass and retained in food webs (Pomeroy et al. 1998; Bardgett et al. 2008; Ducklow 2008). Many factors control how bacteria process carbon, but one of the most important is bacterial growth efficiency (BGE). BGE is the proportion of assimilated organic carbon that is converted into bacterial biomass (del Giorgio and Cole 1998). When BGE is high, more carbon is turned into bacterial biomass where it can be retained for longer periods of time while also serving as a resource for other members of the food web. In contrast, when BGE is low, microbially assimilated carbon has a shorter residence time and is released into the environment as CO_2_. Typically measured as an aggregate property of the microbial community, BGE is notoriously variable among habitats and has proven difficult to predict (del Giorgio and Cole 1998; Sinsabaugh et al. 2013). While a range of chemical and physical variables influence BGE at the community-level (Apple and del Giorgio 2007; Hall and Cotner 2007; del Giorgio and Newell 2012; Geyer et al. 2016), fewer studies have investigated how the traits of species contribute to BGE (Pold et al. 2020).

A trait-based approach provides an opportunity for a deeper understanding of how microbial composition and physiology contribute to BGE. By focusing on physiological, morphological, or behavioral characteristic that affect performance, a trait-based approach can be used to predict fitness under a set of environmental conditions (Lennon et al. 2012). The distribution of traits among organisms may reflect adaptations, phylogenetic relatedness, and metabolic constraints (Martiny et al. 2015). In the context of BGE, insight may be gained by identifying taxon-specific differences in microbial metabolism that result from the physiological balance between cellular growth and energetic demands. For example, bacterial growth strategy is predicted to constrain BGE via physiological trade-offs (Litchman et al. 2015). As a result, it has been hypothesized that oligotrophs have higher maximum growth efficiency than copiotrophs (Roller and Schmidt 2015), and rapidly growing bacteria have been shown to “spill” up to 20 % of their energetic budget due to overflow respiration (Russell 1991, 2007). Likewise, species that specialize on only a few resources are predicted to be more efficient at using those resources than generalist species (Dykhuizen and Davies 1980; Glasser 1984). Together, traits such as maximum growth rate and the number of resources used (i.e., niche breadth) could underlie species-specific differences in BGE.

A trait-based approach to BGE also requires that metabolism be examined with respect to the resources that are being consumed. Different resources can affect cellular ATP yield depending on the metabolic pathways used (Fuhrer et al. 2005; Flamholz et al. 2013), which in turn can influence cellular growth yield (Neijssel and de Mattos 1994; Russell and Cook 1995). For example, glucose is metabolized via glycolysis, but growth on more complex, aromatic compounds, such as protocatechuate, requires the β-ketoadipate pathway which yields less ATP (Gottschalk 1986). Furthermore, energy-producing catabolic processes and biomass-producing anabolic processes are not independent (Russell and Cook 1995; Kempes et al. 2012). For example, cells have the potential to produce > 30 ATP from a single glucose molecule if it is completely oxidized, but there would be no remaining carbon to yield new biomass. Instead, cells must use the intermediate products of glycolysis and the TCA cycle to form proteins and other cellular material, which diminishes the maximum potential ATP yield (Gottschalk 1986; Flamholz et al. 2013). In addition, biomass production requires materials and energy. For example, the synthesis of proteins, which constitute ∼70 % of cellular dry mass, requires amino acid building blocks and 4 ATP per peptide bond (Tempest and Neijssel 1984; Gottschalk 1986; Lynch and Marinov 2015). Therefore, because resources differ in their potential energy yield and bacteria differ in their ability to extract energy and form biomass from a given resource, BGE should vary based on the resources available to bacteria.

In this study we used a trait-based approach to understand how species identity and resource type control BGE. We measured BGE in a diverse set of bacterial isolates supplied with one of three different carbon resources that varied in chemical structure and metabolic pathway (Fig. S1). The trait-based approach provides a framework to understand how and why the composition of microbial communities should affect ecosystem functioning (Wallenstein and Hall 2012; Krause et al. 2014). We used the taxonomic and phylogenetic relatedness of the bacterial isolates to explore the variation in BGE when supplied with different carbon resources. In addition to partitioning variation in BGE based on species identity and resource type, we tested for hypothesized trade-offs with growth rate and niche breadth while taking phylogeny into account. Furthermore, using the genomes of each isolate, we evaluated whether metabolic pathways could explain differences in BGE among diverse representatives of aquatic bacteria from north temperate lakes. Last, to test if resource type affects the metabolic traits that underlie BGE (i.e., production and respiration), we tested for resource-specific relationships between respiration and production rates for each resource. Together, our trait-based approach provides a framework for understanding linkages between community structure and function due to the physiological constraints on BGE and suggest that large changes in community composition or available resources may alter BGE and therefore carbon cycling in predictive ways.

## Results

### Bacterial growth efficiency

Using measures of bacterial productivity (BP) and respiration (BR), we calculated bacterial growth efficiency (BGE) for 20 aquatic bacterial isolates growing on three different resources: glucose, succinate, and protocatechuate (Fig. S1). Isolates were enriched from north temperate lakes, and all belonged to the Proteobacteria phylum with representatives from the Alpha-, Beta-, and Gammaproteobacteria subphyla (Fig. 1, Fig. S2). Across isolates, we found that BGE ranged from <0.01 to 0.32 (Fig. 1). Based on linear mixed-effects models we found that species identity and resource type explained a substantial amount of variation in BGE. Across resources, species identity explained 58 % of the variation in BGE, and 67 % of the variation within a resource type. The taxonomic order of each species explained 20 % of the variation in BGE across all resources, and 28 % of the variation within each resource type. Resource type only explained 8 % of the variation in BGE across all species, but 63 % of the variation within species (see Table S1 for additional information model output).

**Fig. 1:**
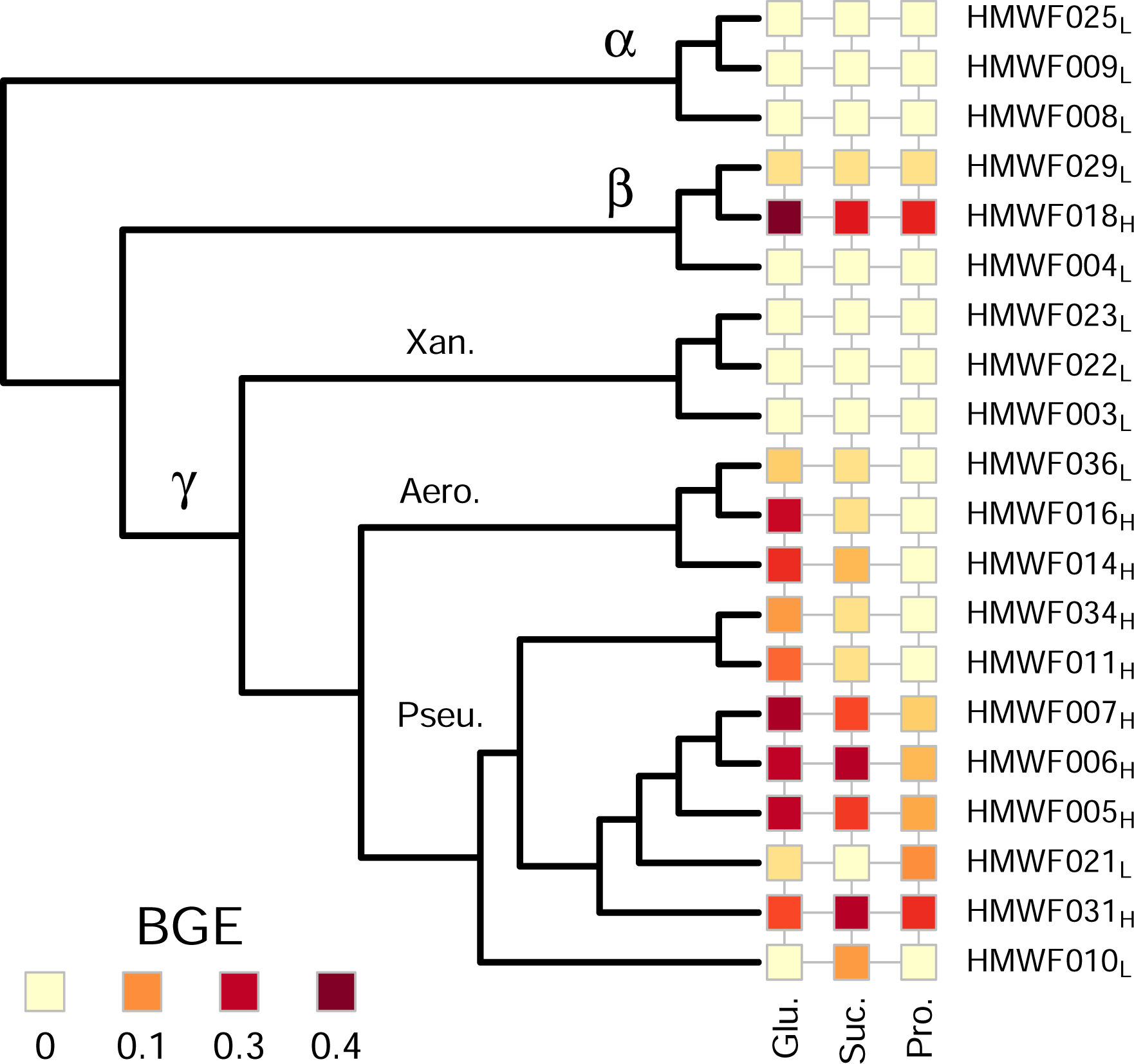
Bacterial growth efficiency (BGE) of each isolate for each resource. BGE was calculated based on measured production (BP) and respiration (BR) rates using the following equation: BGE = BP / (BP + BR). Cladogram is based on the RAxML consensus tree but is shown without branch lengths for visualization (see Fig. S2 for complete phylogenetic tree). Taxonomic class and order are included based on the Ribosomal Database Project taxonomy: *α* = Alphaproteobacteria. *β* = Betaproteobacteria, *γ* = Gammaproteobacteria, *Xan*. = Xanthomonadales, *Aero*. = Aeromondales, *Pseu*. = Pseudomondales. The BGE group is indicated for each isolate (H = high; L = low).

Next, we tested for phylogenetic signal in BGE. Using Blomberg’s K, no phylogenetic signal was detected for BGE when isolates were supplied with succinate (K = 0.002, *p* = 0.24) or protocatechuate (K = 0.001, *p* = 0.146), but there was a significant phylogenetic signal when isolates used glucose (K = 0.002, *p* = 0.04). However, the low K value suggests that BGE is over-dispersed (i.e., less phylogenetic signal than expected under Brownian motion). Similarly, when we use Pagel’s λ, we found no evidence that BGE had a phylogenetic signal when the isolates were supplied with any of the resources (Glucose: λ = 0.10, *p* = 0.76; Succinate: λ = 0.13, *p* = 0.66; Protocatechuate: λ < 0.01, *p* = 0.99).

Last, we determined if the values of BGE observed across isolates and resources were unimodally distributed. Based on Hartigan’s dip test, we found that there was a bimodal distribution of BGE among our isolates when supplied with glucose or succinate (D_glu_ = 0.07, *p* = 0.58; D_suc_ = 0.08, *p* = 0.30; Fig. S3). Using this distribution, we split isolates into two groups (based on the glucose BGE), which we define as the “high-BGE” (19.6 % ± 2.8), and “low-BGE” (0.5 % ± 0.2) groups.

### Phenotypic comparisons

Using linear models, we identified phenotypic differences between isolates that were related to BGE (Fig. 2). While there was no relationship between BGE and maximum growth rate in the low-BGE group of bacteria (µ_max_; F_1,7_ = 0.035, r^2^ < 0.01, *p* = 0.86), we identified a significant inverse relationship between BGE and µ_max_ for the high-BGE group (F_1,7_ = 7.79, r^2^ = 0.53, *p* = 0.027). This model predicts a 2.6 % decrease in BGE for each per minute increase in µ_max_ in the high-BGE group. In contrast to our predictions, there was no relationship between niche breadth (Levin’s Index) and BGE for the low-BGE group (F_1,7_ = 1.42, r^2^ = 0.17, *p* = 0.27) or the high-BGE group (F_1,7_ = 0.92, r^2^ = 0.11, *p* = 0.37).

**Fig. 2:**
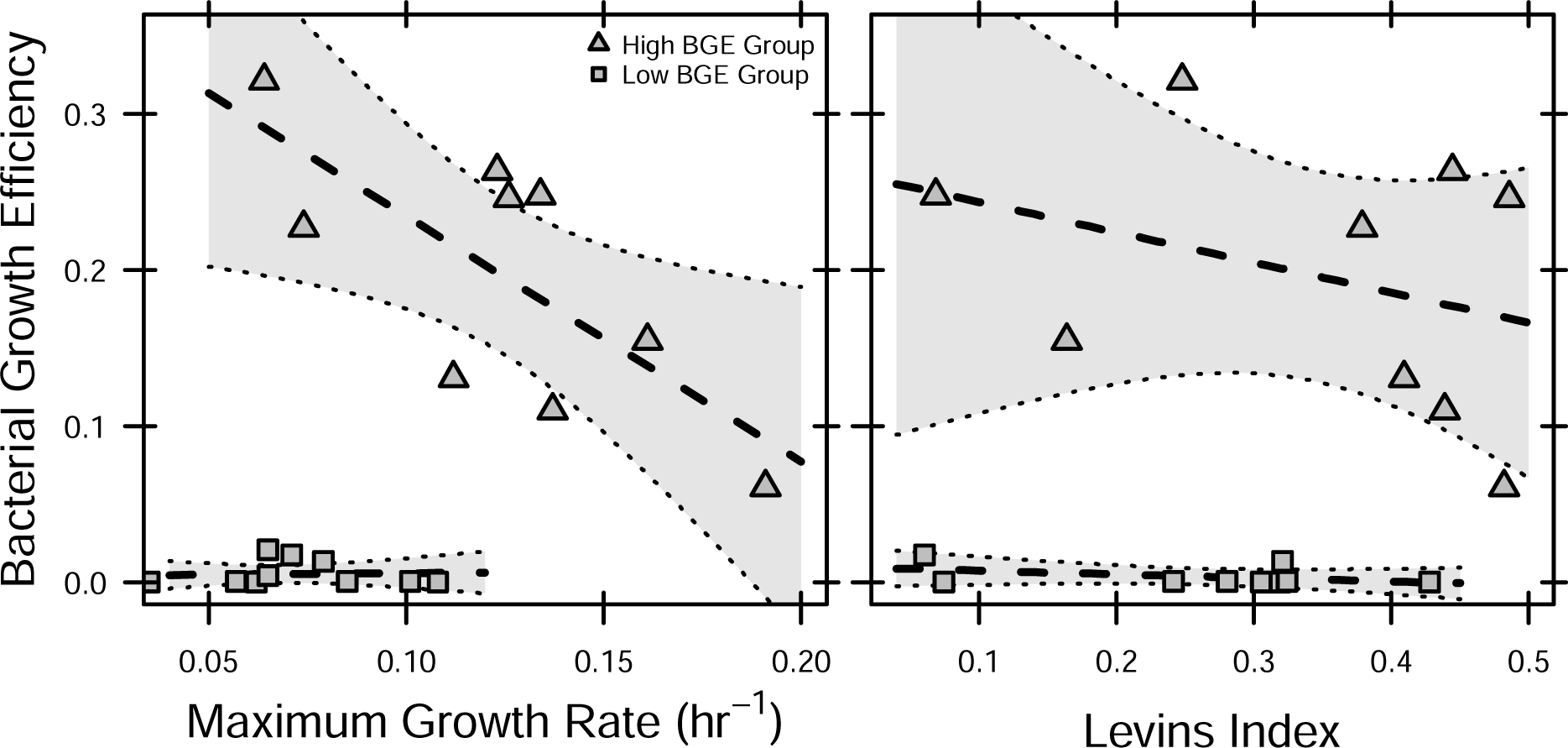
Phenotypic traits associated with BGE. A: Maximum growth rate, a measure of growth strategy, demonstrates a trade-off (negative relationship) with BGE in the high-BGE group (F_1,7_ = 9.52, r^2^ = 0.54, *p* = 0.015), but not the low-BGE group (F_1,7_ = 0.51, r^2^ = 0.06, *p* = 0.50). B: Levin’s Index, a measure of niche breadth, does not demonstrate a trade-off with BGE in either the high or low BGE groups (high: F_1,7_ = 0.92, r^2^ = 0.11, *p* = 0.37; low: F_1,7_ = 1.47, r^2^ = 0.17, *p* = 0.27). High- and low-BGE groups were determined based on the bimodal distribution of BGE (see Fig. S3).

### Genomic comparisons

We detected genomic differences related to BGE. First, isolates in the high-BGE group had 26 % more annotated metabolic pathways (based on an 80 % module completion ratio cutoff) than isolates in the low-BGE group (high = 66 ± 3, low = 52 ± 4, *t*-test: t_18_ = -2.64, *p* = 0.02). Second, we found that the number of metabolic pathways corresponded with BGE when supplied with glucose, but the direction of the relationship depended on the BGE group. For the high-BGE group there was a negative relationship between BGE and the number of pathways (β = -0.006 ± 0.002, r^2^ = 0.48, *p* = 0.04), but for the low-BGE group there was a positive relationship (β = 0.0003 ± 0.0001, r^2^ = 0.37, *p* = 0.05). Next, we found that differences in the metabolic pathway composition could help explain which BGE group an isolate belongs, and the pathway composition of an isolate was related to its BGE. Specifically, we found three pathways that were indicators of an isolate being in the high-BGE group (Table 1). Likewise, for the high-BGE group we found that we could explain 24 % of the variation in BGE based on the composition of metabolic pathways (dbRDA: F_1,7_ = 2.17, R^2^ = 0.24, *p* = 0.05). We found eight pathways with significant positive or negative correlations (|ρ| > 0.7) with BGE (Table 2). However, we did not find a relationship between pathway composition and BGE for the low-BGE group (*p* = 0.45), or evidence that BGE group alone could predict pathway composition (PERMANOVA: *p* = 0.23).

**Table 1:** Genetic pathways unique to the high BGE isolates. Functional metabolic pathways were identified from genome sequencing and predicted using Maple. Prob. = probability statistic from indicator species analysis: the probability that the “species” (i.e., pathway) is not unique to the group.

| Pathway | Reference Function | Prob. |
| --- | --- | --- |
| M00045 | Histidine degradation<br>(histidine → N-formiminoglutamate → glutamate) | 0.02 |
| M00060 | Lipopolysaccharide biosynthesis<br>(Kdo2-lipid A biosynthesis) | 0.02 |
| M00565 | Trehalose biosynthesis<br>(D-glucose-1P → trehalose) | 0.03 |

**Table 2:** Genetic pathways correlated with BGE in the high-BGE group. Correlations are spearman’s rank correlations between BGE and the pathway presence. Pathways with correlation coefficients ≥ |0.70| were considered significant.

| Relationship | $\rho$ | Pathway | Reference Function |
| --- | --- | --- | --- |
| Positive | 0.82 | M00025 | Tyrosine biosynthesis (chorismate → tyrosine) |
|  | 0.72 | M00034 | Methionine salvage pathway |
| Negative | -0.73 | M00117 | Ubiquinone biosynthesis (chorismate → ubiquinone) |
| | -0.82 | M00044 | Tyrosine degradation (tyrosine → homogentisate) |
| | -0.82 | M00053 | Pyrimidine deoxyribonucleotide biosynthesis<br>(CDP/CTP → dCDP/dCTP, dTDP/dTTP) |
| | -0.82 | M00549 | Nucleotide sugar biosynthesis (glucose → UDP-glucose) |
| | -0.82 | M00568 | Catechol ortho-cleavage (catechol → 3-oxoadipate) |
| | -0.82 | M00637 | Anthranilate degradation (anthranilate → catechol) |

### Resource effects

Indicator variable linear regression revealed a positive relationship between the per cell respiration and production rates (Fig. 3, F_9,42_ = 8.07, R^2^ = 0.63, *p* < 0.001) with there being a higher y-intercept for the high-BGE group of isolates (β_Group_ = 2.7, *p* < 0.001). Resource type, however, had no effect on the BR-BP relationship or the effect of BGE group (i.e., no interactions; all *p* > 0.25). Last, we did not find evidence that the slope of the BR-BP relationship was different from one (*t*_42_ = 0.76, *p* = 0.45; Fig. 3); therefore, the two measures of bacterial metabolism scale proportionately (i.e., isometrically) with one another.

**Fig. 3:**
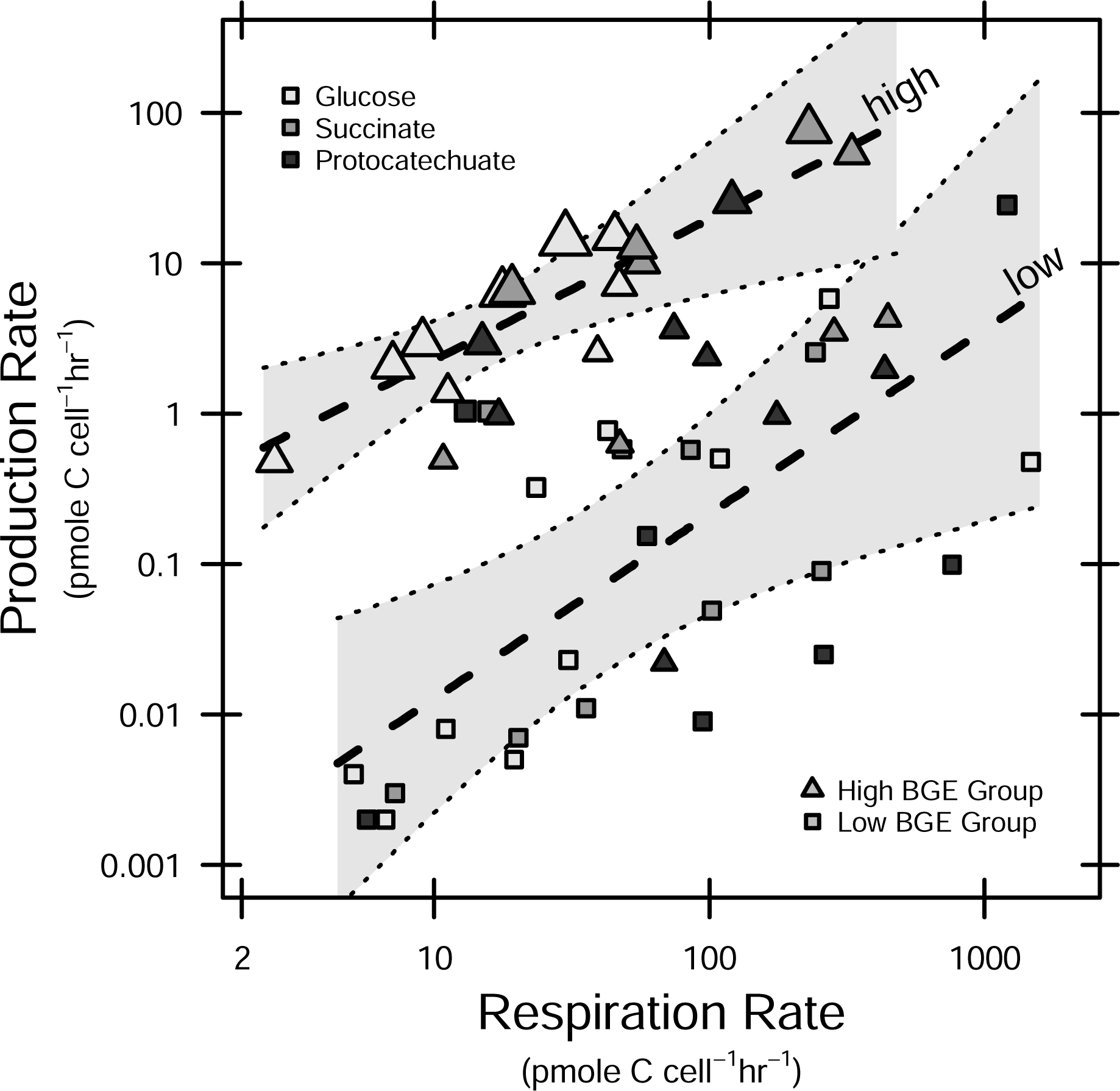
Relationship between respiration and production rates. Respiration and production rates were compared using an indicator variable linear regression (F_9,42_ = 4.92, R^2^ = 0.51, *p* < 0.001). According to the regression model, production rate increases with respiration rate proportionally (i.e., isometric scaling: slope not significantly different from one, *t*_42_ = -0.26, *p* = 0.79). In addition, group (high vs. low BGE) was a significant factor and isolates in the high BGE group had a greater y-intercept (*p* = 0.02). The two regression lines represent the linear fits for the high- and low-BGE groups. Symbols indicates isolate group (high and low BGE), and color indicates the resource being used. Symbol size is scaled by growth efficiency.

## Discussion

We measured bacterial growth efficiency (BGE) in 20 aquatic bacterial isolates supplied with one of three resources that varied in their bioavailability, structure, and pathways required for metabolism (Fig. S1). While BGE varied across isolates, there was no evidence for a strong phylogenetic signal in efficiency. However, a substantial amount (20 %) of the variation in BGE could be explained by an isolate’s taxonomic order while a much smaller amount of the variation (8 %) could be attributed to the particular resource used (Fig. 2). We found evidence for a predicted trade-off between maximum growth rate and efficiency, but only on the most labile resource (glucose) (Fig. 2). Even though it explained 63 % of the variation in BGE within an isolate, resource type did not alter the relationship between respiration and production rate observed across isolates (Fig. 3). Together, we propose that BGE is a complex physiological trait, but resource characteristics may modify species-specific physiological performances. Taxonomic groups of bacteria may have fundamentally different growth efficiencies such that changes in community composition may alter the fate of carbon resources (i.e., biomass versus CO_2_) within the ecosystem (Treseder et al. 2011; Wallenstein and Hall 2012).

### Bacterial growth efficiency as a trait

Our results indicate that there are species-specific properties regulating BGE, which may be conserved at higher taxonomic levels. This conclusion is consistent with the view that BGE represents a complex bacterial trait (i.e., aggregate property of numerous cellular functions) with ecological significance, and that different groups of bacteria have different strategies for carbon allocation. Our phylogenetic analyses suggest that BGE may be an over-dispersed trait (at least with glucose) such that the efficiencies of closely related bacteria are less similar than expected. Though some traits such as phosphorus acquisition, photosynthesis, and methanogenesis are phylogenetically conserved deep in the microbial tree of life (Martiny et al. 2006, 2013), others such as complex carbon metabolism are not (Zimmerman et al. 2013). Therefore, BGE may be similar to traits such as complex carbon metabolism that are not deeply conserved. An alternative explanation for this pattern is that our culture collection lacked phylogenetic resolution within some of our taxonomic groups (e.g., Betaproteobacteria) or that the variation in BGE within a taxonomic group (e.g., order) may not be the same across taxonomic groups. Likewise, because our analysis focused on Proteobacteria with a large representation from the Gammaproteobacteria, it is possible that we missed important phylogenetic patterns found in other important lineages (e.g., Bacteroidetes). Regardless, our data reveal that BGE is a complex bacterial trait that is influenced by taxonomic affiliation. As such, it may be possible to make predictions about BGE and other ecosystem functions given information about composition of resident microbiomes (Goberna and Verdú 2016).

### Bacterial growth efficiency on different resources

in resource complexity and the metabolic pathways required for degradation may explain species-specific differences in BGE due to the resource type used. Within an isolate, resource type accounted for 63 % of the variation in BGE. Given that different resources are processed via different metabolic pathways, it is perhaps not surprising that we observed resource-based variation in BGE within species. For example, BGE was higher when isolates were supplied with glucose compared to when they were supplied with protocatechuate. Glucose is a simple sugar that is able to be metabolized by numerous pathways and converted to acetyl-CoA (Neidhardt 2007). Protocatechuate, on the other hand, is a complex aromatic compound that requires a specific metabolic pathway to be converted to acetyl-CoA. Furthermore, because protocatechuate is chemically more complex, it requires more energy (i.e., ATP) to be degraded than more labile resources such as glucose (Harwood and Parales 1996). Therefore, resource complexity and the metabolic pathways required may explain the within-species variation in BGE. Across species, we did not find resource-specific differences in the relationship between respiration and production rate. However, we recognize that our results may be limited by the number and types of resources used in this study. Regardless, our findings suggest that the energetic demands required to use different resources may be a species-specific trait. That is, the energetic demands for individual species may be constrained and therefore not change much when growing on different resources. Together, these findings suggest that the effect of resources on the efficiency of entire microbiomes may depend on the composition of bacteria consuming those resources.

### Bacterial growth efficiency groups

Although, the range of BGE measured across isolates is similar to the range observed in many ecosystems (del Giorgio and Cole 1998), our results suggest that some species of bacteria grow relatively inefficiently, irrespective of resource quality. Across all isolates, we found a bimodal distribution of BGE suggesting that there were two distinct groups with contrasting efficiencies. One group had low BGE (< 5 %) across all resource types while the other group ranged in BGE from 7-30 % (Fig. 1 & 2). One explanation is that the minimum cellular energetic demand (i.e., cellular maintenance costs) is higher in some bacteria than others (Russell and Cook 1995), however, this would likely only have an impact when growth rates are low. Furthermore, energetic demand may be higher when bacteria are grown in minimal media where they must produce all cellular components from a single carbon resource (Tao et al. 1999). Alternatively, nutrient concentrations (e.g., phosphorus) and other physical properties (e.g., temperature) may regulate efficiency (Smith and Prairie 2004; Frey et al. 2013) and the effects of these properties may be species-specific. As such, it is possible that maintenance costs, resource imbalances, and the physical growth conditions affected BGE of our isolates. Furthermore, differences in low-BGE and high-BGE isolates was also reflected in genomic content, including the number and presence-absence of metabolic pathways. However, these genomic features seem to best explain large-scale rather than fine-scale differences in BGE. Together, these findings suggest that there are fundamental differences between bacterial species that determine BGE.

### Physiological trade-offs

We found evidence to support a trade-off between maximum growth rate and BGE (Fig. 2), which is predicted in microbial and non-microbial systems (Glasser 1984; Roller and Schmidt 2015). For example, theoretical models of microbial communities predict a rate-efficiency trade-off (Allison 2014), which has been observed across microbial taxa (Lipson 2015). Physiologically, the trade-off is based on allocation constraints imposed by the balance between energy requirements and biomass yield: organisms with higher maximum growth rates may have greater energetic requirements and thus lower BGE than organisms with lower maximum growth rates (Russell and Baldwin 1979; Russell and Cook 1995). Furthermore, processes that limit respiration, such as oxygen availability, have been shown to suppress bacterial growth rate (Meyenburg and Andersen 1980). Therefore, respiration rate is likely a major control on biomass production and BGE. Consistent with this, we observed an isometric scaling relationship between respiration and production rates (Fig. 3). The non-zero intercept of this relationship suggests that there is a minimum respiration rate required before any biomass can be produced, which is commonly interpreted as the cellular maintenance requirement. Therefore, it is possible that the maintenance energy demand of a bacterial species explains the physiological trade-off between maximum growth rate and growth efficiency.

Theory also predicts a trade-off between resource niche-breadth and growth efficiency (Glasser 1984). This trade-off is based on the assumption that there is an energetic cost to maintaining numerous metabolic pathways (Johnson et al. 2012). As such, species with more metabolic pathways should have more energetic requirements and thus lower BGE; although, the effects of genome reduction has been debated (Giovannoni et al. 2005; Livermore et al. 2014). In this study, we did not find evidence of a trade-off between resource niche breadth and BGE (Fig. 2). Likewise, we did not find evidence that the number of genes or genome size directly influenced BGE (Table S2–S3), but we did find an inverse relationship between the number of pathways and BGE for the high-BGE group. One possible explanation is that the resources used in our phenotypic assay (i.e., Ecolog plates) did not reflect the full metabolic potential of our isolates. Alternatively, there may not be a strong trade-off between niche breadth and efficiency, but further experiments with additional isolates and resources are required to test this prediction more rigorously.

### Genomic signatures

In addition to the physiological differences documented among our isolates, we found genomic evidence of metabolic pathways that are associated with BGE. Specifically, we detected genomic differences between isolates that belong to low-BGE and high-BGE groups. Isolates from the high-BGE group had 26 % more annotated metabolic pathways than isolates in the low-BGE group. Furthermore, we identified three pathways that were unique to the high-BGE group (Table 1) and a number of pathways that were correlated with the observed BGE (Table 2; for more information see Table S2). Together, our findings suggest that there are genomic features that may contribute to or regulate BGE.

In general, the genomic composition of BGE groups appear to reflect differences in cellular biosynthesis. It is possible that species with particular biosynthesis pathways may generate essential cellular components with less energetic demand. For example, the low-BGE isolates lacked some metabolic pathways, including pyridoxal biosynthesis and histidine degradation, which were present in the high-BGE group. The pyridoxal biosynthesis pathway produces vitamin B_6_ from erythrose-4-phosphate (Mukherjee et al. 2011). Because vitamin B_6_ is essential for growth, the isolates lacking the pyridoxal pathway must use alternatives such as uptake from the environment if they are auxotrophic (i.e., unable to synthesize) or other synthesis pathways such as the deoxyxylulose-5-phosphate synthase (DXS) pathway (found in all but three of the genomes in this study; Table S4; Mukherjee et al. 2011). However, the DXS pathway requires pyruvate (a precursor for Krebs cycle) and thus may limit central metabolism and possibly lead to lower BGE. Likewise, the histidine degradation pathway is used to breakdown histidine into ammonium and glutamate (Bender 2012). Alternatively, glutamate can by synthesized from α-ketoglutarate; however, because α-ketoglutarate is an intermediate component of Krebs cycle this may limit central metabolism and possibly lead to reduced BGE if the energetic requirements are maintained but the ability to recycle biomass is reduced.

## Conclusion

A trait-based approach can provide a mechanistic link between the structure and function of bacterial communities. At the cellular level, BGE reflects the balancing of energetic and cellular growth demands. We found evidence of this based on physiological trade-offs (i.e., maximum growth rate) as well as metabolic pathways. As such, changes in community composition and resource availability have the potential to alter food web and ecosystem function due to changes in BGE. For example, communities dominated by species with low BGE should yield a higher net release of CO_2_ from the ecosystem. Alternatively, communities comprised of individuals with high BGE should yield a net increase in ecosystem productivity. However, variation in BGE can arise within a species due to the ways in which it processes different resources. Therefore, changes in the resource supply will alter the performance of individual taxa, but we predict that these changes will not be as strong as changes in BGE that arise owing to differences in community composition. Our results highlight how bottom-up, trait-based approaches may be useful for understanding complex microbial communities in nature.

## Methods

### Bacterial isolates

Using a novel cultivation approach, we isolated 20 bacterial strains from lakes in the Huron Mountain Research Preserve (Powell, MI, USA) by incubating inert carbon beads (Bio-Sep Beads) in the water column for one week. Prior to the incubations, the beads were saturated with a sterile, complex-carbon substrate, i.e., Super Hume (CropMaster, United Agricultural Services of America, Lake Panasoffkee, Florida, USA). Super Hume is a lignin-rich resource comprising 17 % humic and 13 % fulvic acids, and has been shown to be an analog of terrestrial DOC in aquatic ecosystems that can be used by diverse bacteria (Lennon et al. 2013). We used this enrichment technique to select for bacteria with a range of metabolic potentials (Ghosh et al. 2009). After the incubation, beads were rolled on R2 agar plates (BD Difco, Sparks Maryland, USA) and incubated at 25 °C. We picked random colonies from plates and serially transferred until axenic. All isolates were preserved in 25 % glycerol at -80 °C.

We identified each bacterial strain by direct sequencing the 16S rRNA gene. We obtained genomic DNA from log phase cultures using the FastPrep DNA extraction kit according to the manufacturer’s specifications (MP Biomedical). We used 10 ng of genomic DNA to amplify the 16S rRNA gene using the 27F and 1492R bacterial primers (See for primer sequences and PCR conditions). We sequenced the PCR products at the Indiana Molecular Biology Institute (IMBI) at Indiana University (Bloomington, Indiana, USA). Raw sequence reads were quality-trimmed based on a Phred quality score of 25. Forward and reverse reads were manually merged after aligning sequences to the Silva 16S SSU rRNA reference alignment (release 132) using SINA v. 1.2.11 and the Bacteria variability profile (Pruesse 2011). After merging into full length 16S rRNA sequences, we used mothur (Schloss et al. 2009) to check the quality of sequences and alignments were checked using the ARB software package (Ludwig et al. 2004). Finally, sequences were compared to the Silva All-Species Living Tree Project database (Yilmaz et al. 2014) for taxonomic identification (Fig. S2).

### Bacterial growth efficiency

We measured BGE for each isolate when supplied with one of three different carbon substrates: glucose, succinate, or protocatechuate (Fig. S1). These carbon sources (i.e., resources) were chosen based on differences in their bioavailability and structure but also the required pathways for metabolism (see Fig. S1). We measured bacterial respiration and production rates and then calculated BGE as BP/(BP + BR), where BP is bacterial productivity and BR is bacterial respiration (del Giorgio and Cole 1998). BP and BR were measured using triplicate cultures of each isolate. Cultures of each isolate were grown in R2 broth (BD Difco, Sparks Maryland, USA) until mid-log phase. We then transferred 100 µL of culture into 10 mL of M9 broth (Green and Sambrook 2012) with the appropriate carbon source (25 mM C) and allowed 24 h for the cultures to acclimate. We then transferred 100 µL of culture into 10 mL of fresh carbon-amended M9 broth and incubated 1-3 h to replenish nutrients. Using these transfers, we were able to establish populations of each isolate at target cell densities between 10^4^ and 10^5^ cells mL^−1^. We used the populations to measure BP and BR, which were normalized to cell density using plate counts of colony forming units. We measured BP using the ^3^H-Leucine assay (Smith and Azam 1992) with 1.5 mL of culture. We added ^3^H-Leucine to a final concentration of 50 nM and incubated for 1 h. Following incubation, we terminated production with trichloroacetic acid (final concentration 3 mM) and measured leucine incorporation using a liquid scintillation counter. We measured BR using an automated O_2_ measurement system (PreSens Sensor Dish System, PreSens, Regensburg, Germany) on 5 mL of culture based on the rate O_2_ consumption during three-hour incubations. We estimated BR as the slope of O_2_ concentration during the incubation using linear regression. We used a respiratory quotient conversion factor to convert O_2_ depletion into C respiration assuming aerobic growth (del Giorgio and Cole 1998).

### Taxonomic and phylogenetic relationships

We compared differences in BGE across isolates and resources using linear models. First, we used a taxonomic framework to compare BGE between isolates (Lennon et al. 2012). Isolates were classified into taxonomic groups based on the species tree constructed in ARB. We then used mixed linear models to compare BGE across taxonomic groups and resources. To test the hypothesis that taxonomy (i.e., at the order level) affects BGE, we nested resource type within isolate in the linear model. To test the hypothesis that the specific resource used affects BGE, we nested isolate within resource type. We identified the best statistical models based on the variation explained (R^2^) and AIC values. Second, we tested if phylogenetic relationships between isolates explained differences in BGE across isolates. We created a phylogenetic tree based on the full-length 16S rRNA gene sequences. We aligned sequences using the SINA aligner (Pruesse et al. 2012) and checked alignments using ARB. We generated a phylogenetic tree using the CIPRES science gateway (Miller et al. 2010). The phylogenetic tree was created using RAxML v.8.2.12 (Stamatakis 2006). We used the GTRGAMMA DNA substitution model and the rapid hill-climbing algorithm to build our maximum likelihood trees, and we used the extended majority rule to find the consensus tree. We used Blomberg’s K and Pagel’s Lambda to compare trait variation (as a continuous variable) across the tree and test if phylogenetic relationships between isolates could explain differences in traits (Pagel 1999; Blomberg et al. 2003). Blomberg’s K is a test for phylogenetic signal that determines if trait variation is better explained by phylogenetic relationships or Brownian motion. Pagel’s Lambda is a test of phylogenetic signal that determines if trait variation differs from Brownian motion. Last, to determine if the distribution of BGE across isolates was unimodal, we used Hartigan’s dip test for unimodality (Hartigan and Hartigan 1985). Hartigan’s dip test is used to determine if a distribution is unimodal by testing the null hypothesis that there is a dip in the distribution. A significant Hartigan’s dip test would suggest that the distribution is unimodal. Alternatively, the distribution has an internal “dip” (reported as D). All statistical tests were conducted in the R statistical environment (R Core Development Team 2013). We used the *nlme* package (Pinheiro and Bates 2011) for the mixed-effects linear models, the *picante* package (Kembel et al. 2015) for the phylogenetic methods, and the *diptest* package (Maechler 2015) for Hartigan’s dip test.

### Phenotypic comparisons and trade-offs

To test the hypothesis that phenotypic differences and physiological trade-offs underlie BGE variation, we compared the maximum growth rate (μmax) and niche breadth of each isolate. First, to test whether BGE was affected by growth strategy (i.e., copiotrophs vs. oligotrophs), we measured the maximum growth rate of each isolate. Bacterial growth rates were measured based on changes in optical density during 18-h incubations. Bacterial isolates were grown in R2 broth in 48-well plates. We incubated plates with continuous shaking and measured optical density every 15 min using a plate reader (BioTek Synergy MX). Growth curves were analyzed by fitting a modified Gompertz growth model (Zwietering et al. 1990; Lennon 2007) to the observed growth curves using maximum likelihood fitting. We used the model fit as our estimate of µmax.

Second, to test whether BGE was affected by niche breadth, we generated carbon usage profiles using BioLog EcoPlates™ (Garland and Mills 1991). The EcoPlate is a phenotypic profiling tool consisting of 31 unique carbon sources. In addition to the carbon source, each well contains a tetrazolium dye, which in the presence of NADH is reduced resulting in a color change. We used this colorimetric assay to generate carbon usage profiles for each strain. We standardized profiles for each strain by subtracting water blanks (average water blank + 1 SD), and relativizing across substrates. Using these data, we calculated resource niche breadth using Levins Index (Colwell and Futuyma 1971).

We used an indicator variable linear regression to test for changes in BGE rate due to maximum growth rate and niche breadth. We included the BGE group (high-versus low-BGE) as the categorical predictor and BGE as the continuous predictor (Lennon and Pfaff 2005). In addition, we included the interactions term. Where the interaction term was significant, we report the main effects of each categorical predictor (i.e., BGE group). All statistical tests were conducted in the R statistical environment.

### Genomic comparisons

To test the hypothesis that variation in metabolic pathways could explain differences in BGE, we compared the genomes of each isolate. First, we determined the metabolic pathways found in the genome of each isolate. We characterized each isolate using whole genome sequencing. Genomic DNA libraries for each isolate were prepared using the Illumina TruSeq DNA sample prep kit using an insert size of 250 base pairs (bp). Libraries were sequenced on an Illimina HiSeq 2500 (Illumina, San Diego, GA) using 100-bp paired-end reads at the Michigan State University Research Technology Support Facility. We processed raw sequence reads (FASTQ) by removing the Illumina TruSeq adaptors using Cutadapt (Martin 2011), interleaving reads using Khmer (McDonald and Brown 2013), and quality-filtering based on an average Phred score of 30 using the FASTX-toolkit (Hannon Lab 2010). Finally, we normalized coverage to 25 based on a k-mer size of 25 using Khmer. We assembled the genomes using Velvet (Zerbino and Birney 2008) after optimizing assembly parameters for each isolate with Velvet Optimizer (Gladman and Seemann 2012). We annotated contigs larger than 200 bp using Prokka (Seemann 2014), and predicted metabolic and physiological functions using MAPLE with bidirectional best-hit matches (Takami et al. 2012). We identified functional pathway based on the presence of intermediate genes within a pathway. We scored pathways as functional if more than 80 % of the intermediate genes were recovered in the genomes based on module completion ratios.

To test the hypothesis that metabolic pathways affect BGE, we used multivariate methods to compare the pathways of each isolate. First, we used PERMANOVA to determine if there were differences in pathways associated with different levels of BGE, and we used indicator species analysis (Dufrene and Legendre 1997) to determine which metabolic pathways contributed to group differences in BGE. Next, to determine if metabolic pathways could explain differences in BGE within a group, we used distance-based redundancy analysis (dbRDA) which is a multivariate technique that tests if a quantitative predictor can explain differences in multivariate datasets (Legendre and Legendre 2012). Because we scored pathways as present or absent, metabolic distances between isolates were calculated using the Jaccard Index. We tested for significance using a permutation test. If the dbRDA model was significant, we used Spearman’s rank-order correlation to test for correlations between BGE and individual metabolic pathways. We used the *vegan* R package (Oksanen et al. 2012) for multivariate analyses.

### Resource effects

To test the hypothesis that resources have different effects on components of metabolism that affect BGE, we used a linear model to test for a relationship between BR and BP. Because BP requires energy through respiration, we used production rate as the dependent variable and respiration rate as the independent variable. We used an indicator variable linear regression to test for changes in BP rate due to BR. We included the specific resource used and group (high-versus low-BGE) as the categorical predictors and BR as the continuous predictor (Lennon and Pfaff 2005). In addition, we included all interactions terms. Respiration and production rates were log_10_-transformed to meet model assumptions. Last, to determine if the relationship between BR and BP rates was isometric (proportional scaling, slope = one) or allometric (disproportional scaling, slope ≠ one), we used a one-sample t-test to determine if the slope was different from one. All statistical tests were conducted in the R statistical environment.

## Supporting information

Supplemental_GenomeStats

Supplemental_Tree

## Acknowledgements

We thank BK Lehmkuhl and MA Carrison for technical assistance and JB McKinlay and members of the Lennon Lab for critical feedback on an earlier version of this manuscript. This work was supported by the Huron Mountain Wildlife Foundation (MEM & JTL), the National Science Foundation (DEB-0842441, DEB-1442246, and DEB-1501164), and US Army Research Office (W911NF-14-1-0411). All code and data used in this study can be found in a public GitHub repository (https://www.github.com/LennonLab/MicrobialCarbonTraits). Isolate genomes are available on NCBI (BioProject PRJNA420393).

