## Supplemental for "Trait-based approach to bacterial growth efficiency"

**Supplemental Methods**

**Primer Sequences**

Forward (27F): AGAGTTTGATCMTGGCTCAG

Reverse (1492R): TACCTTGTTACGACTT

**PCR Conditions**: Concentrations (Conc.) are given per 50 μL reaction (Rxn.). Components are from the Promega Go-Taq Kit.

| **Component** | **Stock Conc.** | **Final Conc** | **µL per 50 µL Rxn.** |
| --- | --- | --- | --- |
| GoTaq Reaction Buffer | 5 X | 1 X | 10 |
| dNTP Mix (10 mM each) | 40 mM | 200 µM | 1 |
| Forward Primer (10 mM) | 10 µM | 0.2 µM | 1 |
| Reverse Primer (10 mM) | 10 µM | 0.2 µM | 1 |
| GoTaq DNA Polymerase | 5 U/µL | 1.25 U | 0.25 |
| Molecular Grade Water |  |  | 35.75 |
| Template DNA | 10 ng/L | 10 ng | 1 |

**Thermal Cycler Conditions:**

| **Temperature (ºC)** | **Time (sec.)** | **Cycles** |
| --- | --- | --- |
| 94 | 180 |  |
| 94 | 45 | 30 |
| 50 | 30 |  |
| 72 | 90 |  |
| 72 | 600 |  |
| 4 | hold |  |

**Supplemental Tables**

**Table S1:** Output from linear mixed-effects models describing effect of taxonomy and resource type on bacterial growth efficiency (BGE). Mixed-effects models were fit by REML using the *lme()* function in the *nlme* R package. In these models there is a main variable and a nested variable. For each analysis, the variation explained by the main variable is accounted for before the variation explain by the nested variable is determined. As such, these results indicate the relative importance of each variable when grouped together in a nested framework.

| Model | Main Variable | Nested Variable | Variance Explained by Isolate | Variance Explain by Resource | AIC |
| --- | --- | --- | --- | --- | --- |
| Isolate Identity | Species | Resource | 58.3 % | 67.1 % | -48.29 |
| Taxonomic Order | Order | Resource | 19.9 % | 27.6 % | -94.25 |
| Resource Type | Resource | Species | 7.99 % | 63.2 % | -116.73 |

**Table S2:** Genome summary for the Huron Mountain Wildlife Foundation (HMWF) culture collection. Total number of genes is the number of annotated genes. Coverage estimate was calculated based on average contig coverage during assembly. MCR = module completion ratio from Maple analysis. For additional info see GenomeStats.xlsx or NCBI PRJNA420393.

| Strain | Genome length (nucleotides) | # of Genes (total) | # of Pathways (MCR > 80) | Estimated Coverage | Scaffold N50 |
| --- | --- | --- | --- | --- | --- |
| HMWF001 | 4,588,649 | 4,420 | 52 | 15x | 6,682 |
| HMWF003 | 4,460,324 | 4,255 | 47 | 13x | 14,493 |
| HMWF004 | 5,476,832 | 6,021 | 41 | 5x | 4,613 |
| HMWF005 | 5,872,459 | 6,493 | 66 | 7x | 5,177 |
| HMWF006 | 6,285,913 | 6,186 | 57 | 7x | 14,895 |
| HMWF007 | 6,051,858 | 6,348 | 63 | 6x | 8,597 |
| HMWF008 | 3,802,708 | 4,018 | 35 | 11x | 7,977 |
| HMWF009 | 3,919,360 | 3,860 | 57 | 14x | 19,491 |
| HMWF010 | 5,890,869 | 6,195 | 66 | 12x | 35,798 |
| HMWF011 | 6,024,040 | 5,819 | 69 | 10x | 30,117 |
| HMWF013 | 4,611,749 | 4,700 | 58 | 9x | 9,462 |
| HMWF014 | 4,397,736 | 4,521 | 63 | 9x | 11,207 |
| HMWF015 | 4,658,495 | 4,602 | 67 | 12x | 22,382 |
| HMWF016 | 4,523,627 | 4,279 | 74 | 21x | 65,781 |
| HMWF017 | 4,461,702 | 4,205 | 75 | 12x | 43,441 |
| HMWF018 | 5,805,605 | 5,820 | 47 | 18x | 15,242 |
| HMWF019 | 6,906,470 | 5,867 | 55 | 15x | 19,218 |
| HMWF021 | 5,954,435 | 5,610 | 66 | 18x | 59,774 |
| HMWF022 | 4,238,619 | 4,868 | 24 | 4x | 3,647 |
| HMWF023 | 4,584,820 | 4,363 | 56 | 14x | 24,346 |
| HMWF025 | 3,960,983 | 3,764 | 51 | 21x | 44,746 |
| HMWF026 | 3,392,253 | 3,424 | 50 | 18x | 10,819 |
| HMWF028 | 4,488,319 | 4,292 | 48 | 13x | 4,397 |
| HMWF029 | 4,650,641 | 4,374 | 61 | 20x | 23,346 |
| HMWF030 | 4,758,486 | 4,961 | 53 | 9x | 4,075 |
| HMWF031 | 10,014,416 | 12,286 | 82 | 4x | 2,539 |
| HMWF032 | 4,391,151 | 4,100 | 62 | 37x | 583,475 |
| HMWF034 | 6,119,028 | 5,738 | 72 | 26x | 110,028 |
| HMWF035 | 4,967,466 | 4,664 | 56 | 2x | 69,493 |
| HMWF036 | 4,455,344 | 4,269 | 65 | 11x | 26,720 |

**Table S3:** DXS presence/absence in the Huron Mountain Wildlife Foundation (HMWF) culture collection genomes. The DXS pathway (deoxyxylulose-5-phosphate synthase) is an alternative pathway for the synthesis of vitamin B_6_. BGE Group: L = Low BGE group; H = High BGE group. Results reported for all HMWF genomes but those used in the current study are indicated.

| Genome | DXS | BGE Group | Used in this study |
| --- | --- | --- | --- |
| HMWF003 | TRUE | L | Yes |
| HMWF004 | TRUE | L | Yes |
| HMWF005 | TRUE | H | Yes |
| HMWF006 | TRUE | H | Yes |
| HMWF007 | TRUE | H | Yes |
| HMWF008 | FALSE | L | Yes |
| HMWF009 | TRUE | L | Yes |
| HMWF010 | TRUE | H | Yes |
| HMWF011 | TRUE | H | Yes |
| HMWF013 | TRUE | NA | No |
| HMWF014 | TRUE | H | Yes |
| HMWF015 | TRUE | NA | No |
| HMWF017 | TRUE |  | No |
| HMWF018 | TRUE | H | Yes |
| HMWF019 | TRUE | NA | No |
| HMWF021 | TRUE | L | Yes |
| HMWF022 | FALSE | L | Yes |
| HMWF023 | TRUE | L | Yes |
| HMWF025 | TRUE | L | Yes |
| HMWF026 | TRUE | NA | No |
| HMWF028 | TRUE | NA | No |
| HMWF029 | FALSE | L | Yes |
| HMWF030 | TRUE | NA | No |
| HMWF031 | TRUE | H | Yes |
| HMWF032 | TRUE | NA | No |
| HMWF034 | TRUE | H | Yes |
| HMWF036 | TRUE | L | Yes |
| HWMF016 | TRUE | H | Yes |

**Supplemental Figures**

**Figure S1:** Carbon resources used to study BGE variation in environmental isolates. **A**: Glucose – the baseline resource used to compare BGE across isolates. Glucose can be degraded by the Embden-Meyerhof-Parnas, pentose phosphate, or Entner-Doudoroff pathway. Ultimately, these pathways produce pyruvate (and then acetyl-CoA), which enters Krebs cycle and is used to produce energy and intermediates for biomass synthesis, when cells are grown aerobically. Alternatively, glucose can be fermented into organic acids (e.g., lactate), but these reactions yield less energy (Gottschalk 1986). **B**: Succinate – a simple organic acid. Succinate is an intermediate of Krebs cycle and thus it does not require previous degradation. Additionally, succinate can be directly used to produce energy via succinate dehydrogenase (Neidhardt 2007). **C**: Protocatecuate – is a complex resource with an aromatic core. Typically, it is degraded to acetyl-CoA and succinyl-CoA via the β-ketoadipate pathway (Harwood and Parales 1996). Protocatechuate is commonly used to study aromatic resource degradation in ecosystems, and the β-ketoadipate pathway is commonly found in bacteria across the phylum Proteobacteria (Buchan et al. 2000).

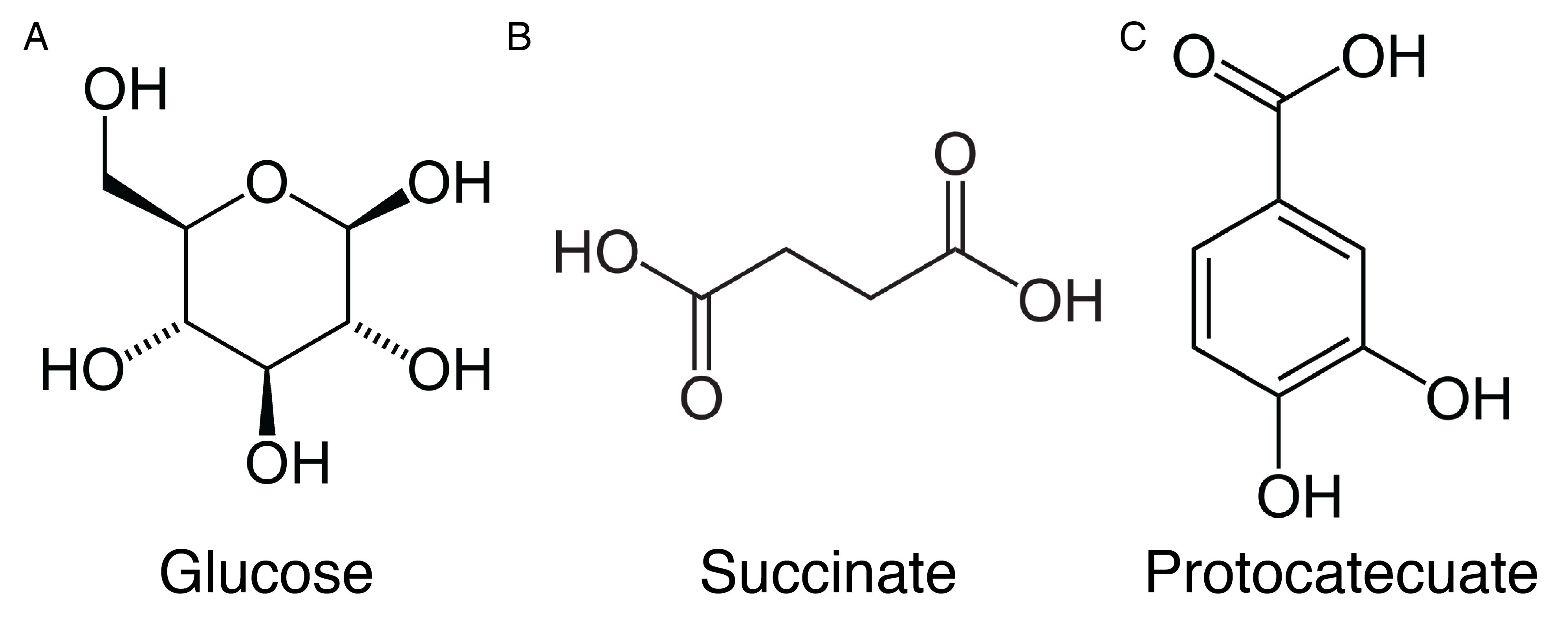

**Figure S2:** Maximum likelihood phylogenetic tree of lake bacterial isolates used to study BGE. Nearest relatives and other type-strains are included as a taxonomic reference. Isolates are organized by taxonomic class. The outgroup (*Aquifex*) is included as the tree root. Scale bar represents 0.01 base substitutions. The RAxML maximum likelihood tree for just the isolates used in the phylogenetic analyses can be found in Supp_tree.nwk.tre (Newick format).

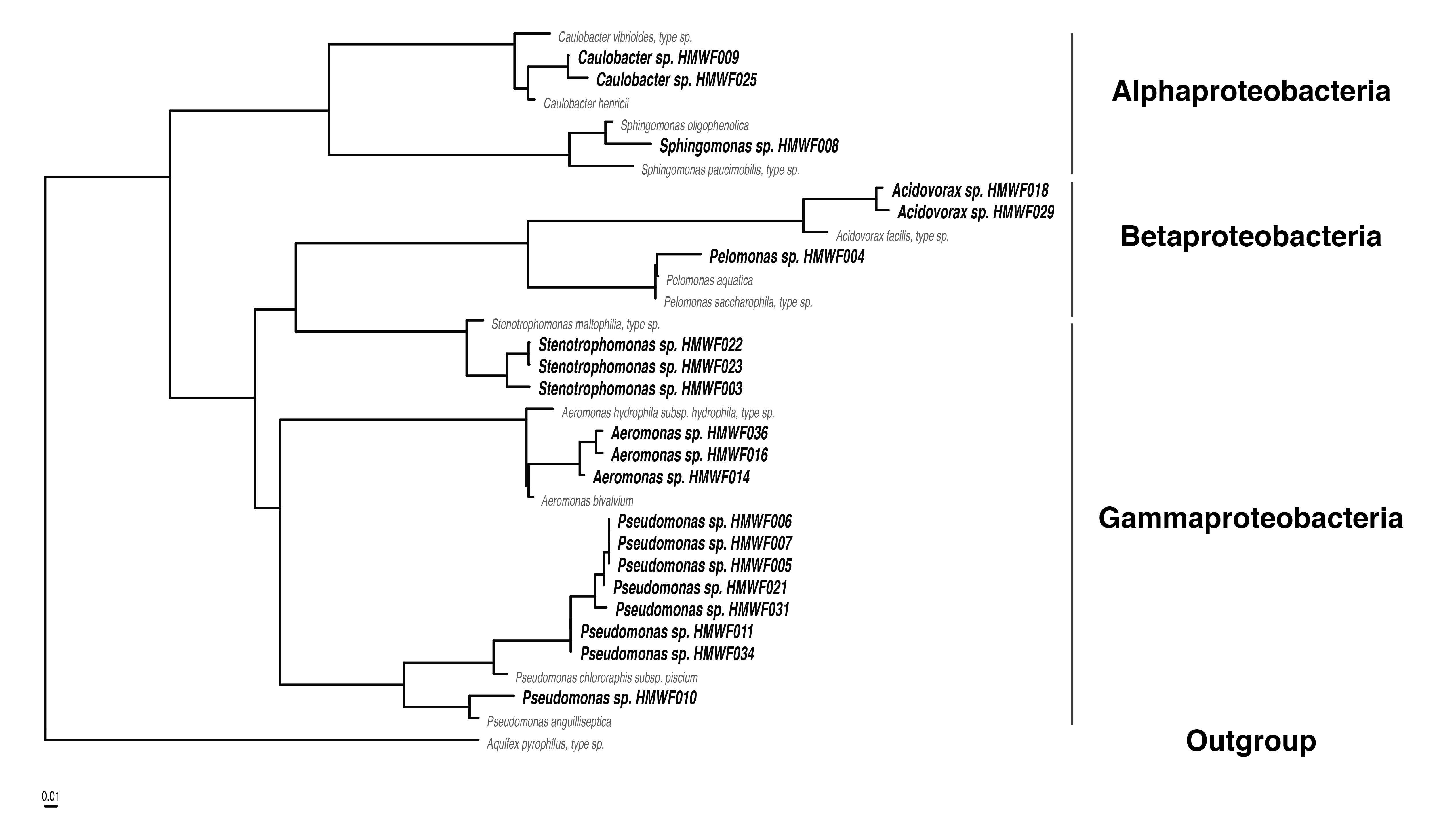

**Figure S3:** Kernel density of BGE when grow on glucose. Based on Hartigan’s dip test, we found that there was a bimodal distribution of BGE among our isolates when supplied with glucose or succinate (D_glu_ = 0.07, *p* = 0.58; D_suc_ = 0.08, *p* = 0.30). High- and low-BGE groups were determined based on the bimodal distribution of BGE.

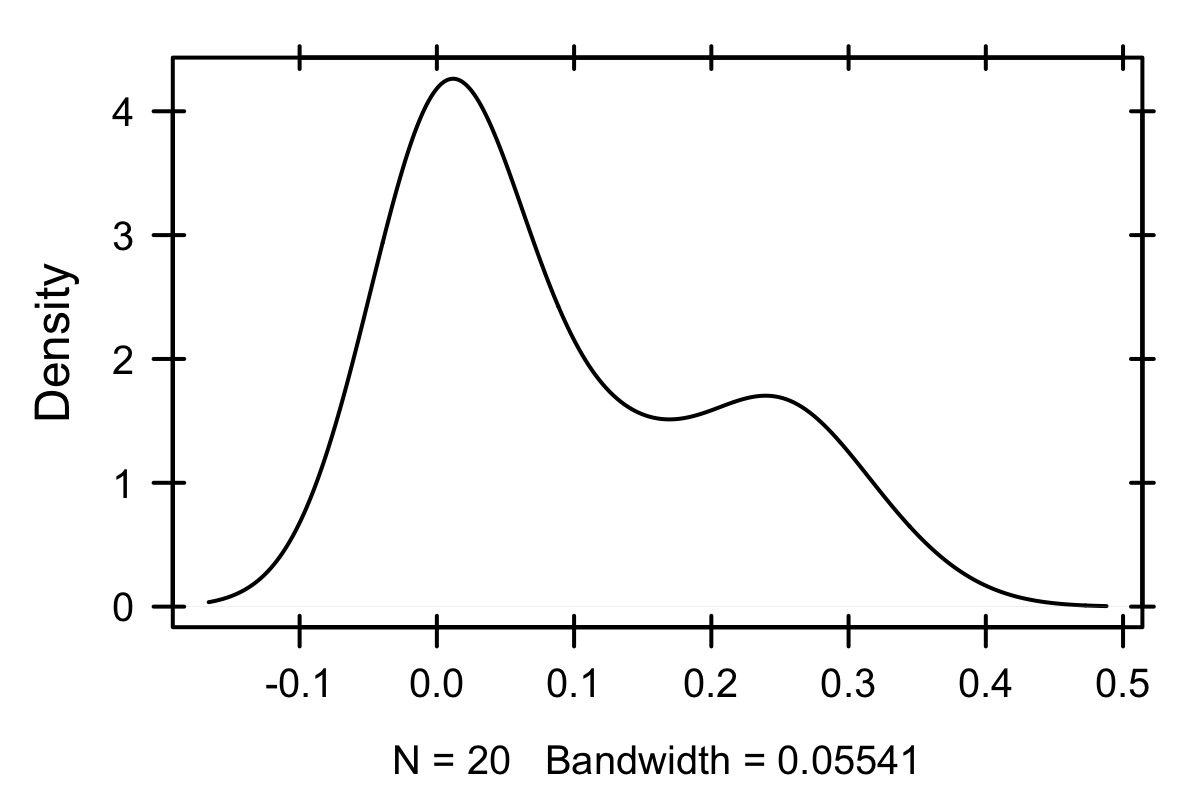
